# scMomentum: Inference of Cell-Type-Specific Regulatory Networks and Energy Landscapes

**DOI:** 10.1101/2020.12.30.424887

**Authors:** Larisa M. Soto, Juan P. Bernal-Tamayo, Robert Lehmann, Subash Balsamy, Xabier Martinez-de-Morentin, Amaia Vilas-Zornoza, Patxi San-Martin, Felipe Prosper, David Gomez-Cabrero, Narsis A. Kiani, Jesper Tegner

**Author notes:** These authors conducted the work at King Abdullah University of Science and Technology but are now part of the Department of Human Genetics at McGill University, Montreal, QC H3A 0C7, Canada.

## Abstract

Recent progress in single-cell genomics has generated multiple tools for cell clustering, annotation, and trajectory inference; yet, inferring their associated regulatory mechanisms is unresolved. Here we present scMomentum, a model-based data-driven formulation to predict gene regulatory networks and energy landscapes from single-cell transcriptomic data without requiring temporal or perturbation experiments. scMomentum provides significant advantages over existing methods with respect to computational efficiency, scalability, network structure, and biological application.

**Availability:** scMomentum is available as a Python package at https://github.com/larisa-msoto/scMomentum.git

---

The utilization of single-cell RNA sequencing (scRNA-seq) technologies in several research contexts, such as the Human Cell Atlas (HCA), has shown that different cell types have characteristic transcriptomic signatures [1,2]. The characterization of cell types from scRNA-seq data has been a constant challenge. Although, it is known that reconstructing gene-regulatory networks (GRNs) is a critical step towards understanding the establishment of cellular identity. The idea of modeling transcriptomes with systems biology approaches has generated numerous methods using bulk measurements [3]. However, remaining hurdles include extensive data requirements and interpretability of the resulting networks. Single-cell genomics offers the opportunity to resolve such challenges, enabling the ability to model developmental progressions with classic paradigms [4], such as the Waddington landscape [5].

Although scRNA-seq profiling produces many samples (cells), thus increasing statistical power, there are inherent limitations such as missing transcripts that complicate the distinction of signal from noise [6]. Several existing methods rely on identifying an accurate time progression of cells (pseudotime) [7,8,9]. Such an ordering limits their applicability to time-series experiments and induces potential error sources when inferring pseudotime on non-temporal datasets. A common strategy is to use perturbation experiments to sample cause-effect events [10,11], but these methods are not scalable to interrogate different cell types *en masse*.

Here we explore the assumption that regulatory signals are specific and similar among cells belonging to the same cell type, as they would be confined to move around within their quasistable attractor. This enables us to use a linear approximation while still accounting for a non-zero velocity in the quasi-stable state. This formulation ensures computational efficiency and scalability. In such a linear model, every gene expressed in a given cell type could affect all the other genes and vice versa. The model takes advantage of RNA velocity [12] as the gene-level outcome of the expression of all the genes regulating it, weighted by a directed interaction matrix (the inferred GRN) and their corresponding degradation rate (Figure 1a). With both the weights and directions in hand, we can estimate an energy function [13] such that individual cellular energies capture their developmental potential. When projecting such energies on a twodimensional grid, one can get a robust estimation of the developmental energy landscape recovered by those cells (Figure 1a).

**Figure 1.**
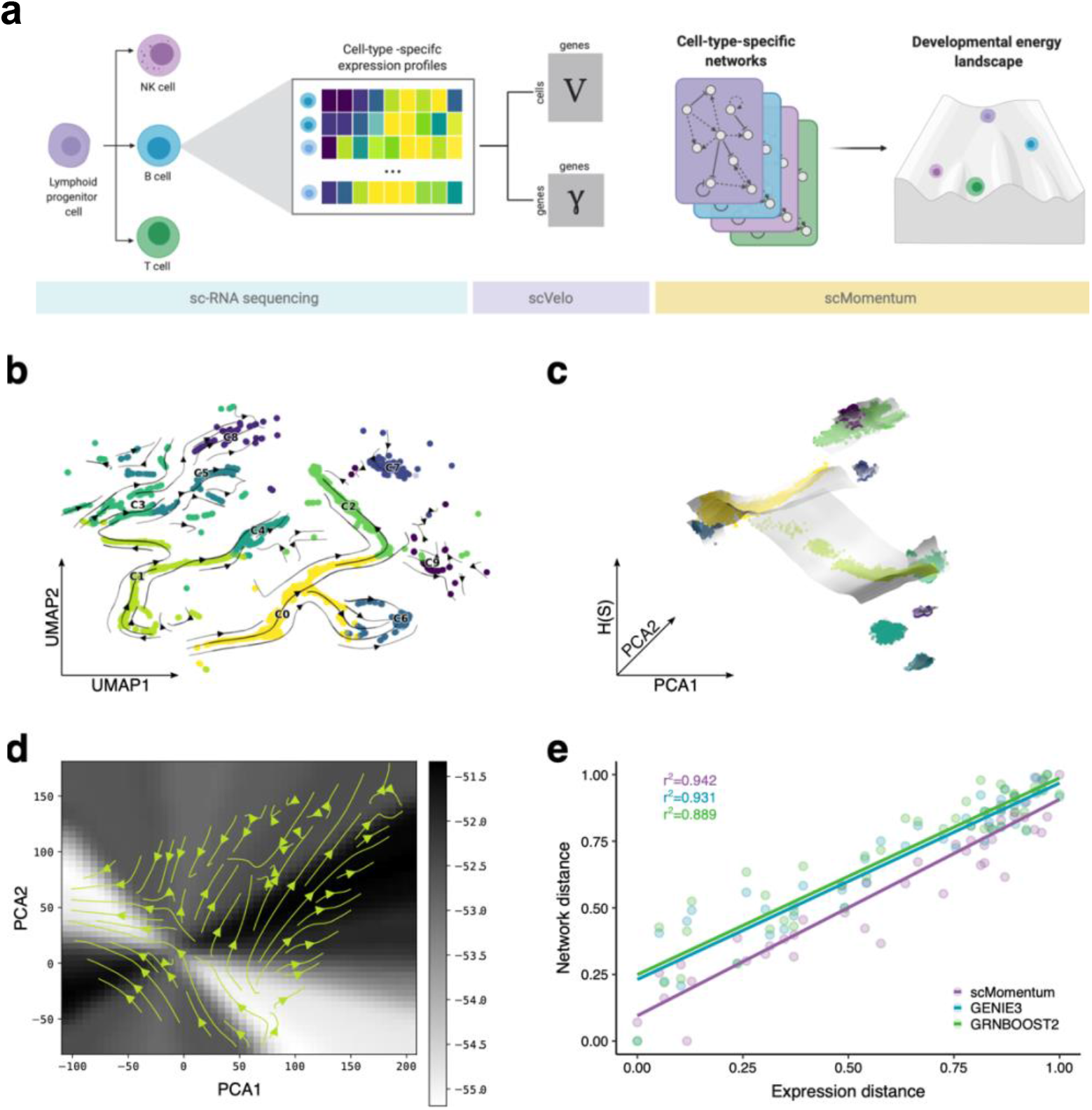
scMomentum captures cluster-specific properties and branching points in a simulated dataset. **(a)** Analysis pipeline upstream of scMomentum. **(b)** Cell-velocity map; cells are colored according to each cluster. **(c)** Energy landscape; cells are colored in the same way as (b); the grayscale of each landscape sheet represents the grid’s energy. **(d)** PCA projection of cell velocities on top of the corresponding energy grid shown in grayscale. **(e)** Methods’ benchmark based on the preservation of cluster distances. Here each point represents the distance between a pair of clusters; r-squared values show the Mantel correlation of expression and network distances for each method, according to the color legend at the bottom right. All correlations were significant at a p-value cut-off of 0.05.

To evaluate this idea, we simulated expression profiles for 500 genes in 20,000 cells using Dyngen [14]. We constructed two independent branching trajectories and used scVelo to estimate RNA velocity and degradation rate (Figure 1b). The inferred GRNs were used to derive the associated energy (Figure 1c). The cells from the branch starting at cluster 1 go towards a lower energy state, and the endpoint clusters are all at a lower energy state than that of their starting point. This agrees with Waddington’s classical proposition that cells will go “*downhill*” on the landscape as they differentiate. On the other hand, the branch starting at cluster 0 follows an upward trend, indicating the need for external inputs to progress along that branch instead of being in a poised state. To further explore the directionality of cells in the landscape, we projected the velocities on top of the energy landscape (Figure 1d). We found a flow from a high to a low energy state. Moreover, the branching is captured both by the velocities and by the energies (Figure 1d).

To assess the performance of scMomentum, we benchmarked it against GENIE3 [15] and GRNOOST2 [16]. Since there is a lack of a widely accepted gold standard, we designed a metric that quantifies the extent of biological signal preservation. We assume that similar cell types have relatively close expression profiles (expression-derived distance). Consequently, if the derived networks are accurate enough, they would also be similar (network-derived distance). Thus, we test the correlation between cell type distance matrices using a Mantel test to account for the matrices’ spatial arrangement (symmetry and row-column relationships). scMomentum showed the highest correlation (Figure 1e), although GENIE3 and GRBOOST2 also had significant correlations in this controlled setting. Thus, we found this to be a reasonable metric to assess the performance of different methods.

We evaluated scMomentum in an in-house-generated human hematopoiesis study and five public data sets (see Methods), totaling more than 200K cells (see supplement for details on the preprocessing steps). We found that the inferred networks preserved the distances between cell types in all the datasets that captured multiple developmental stages at once (dynamic behavior) (Figure 2a). As a negative control, we showed that this was not true for a non-dynamical dataset (mBA18, see Methods), highlighting the need for cellular dynamics when computing RNA velocity and using it to infer GRNs. We benchmarked our approach on the human hematopoiesis dataset and found that scMomentum was better at preserving cell type distances than existing methods (Figure 2b). Although the difference between the coefficients was relatively small, scMomentum holds a significant advantage in network structure and computational efficiency. We could predict GRNs for ~100K cells expressing 1,000 genes in ~1 minute on a machine with a 3.1 GHz Dual-Core Intel Core i5 processor, while GENIE3 and GRNBOOST2 were unable to finish within a day.

**Figure 2.**
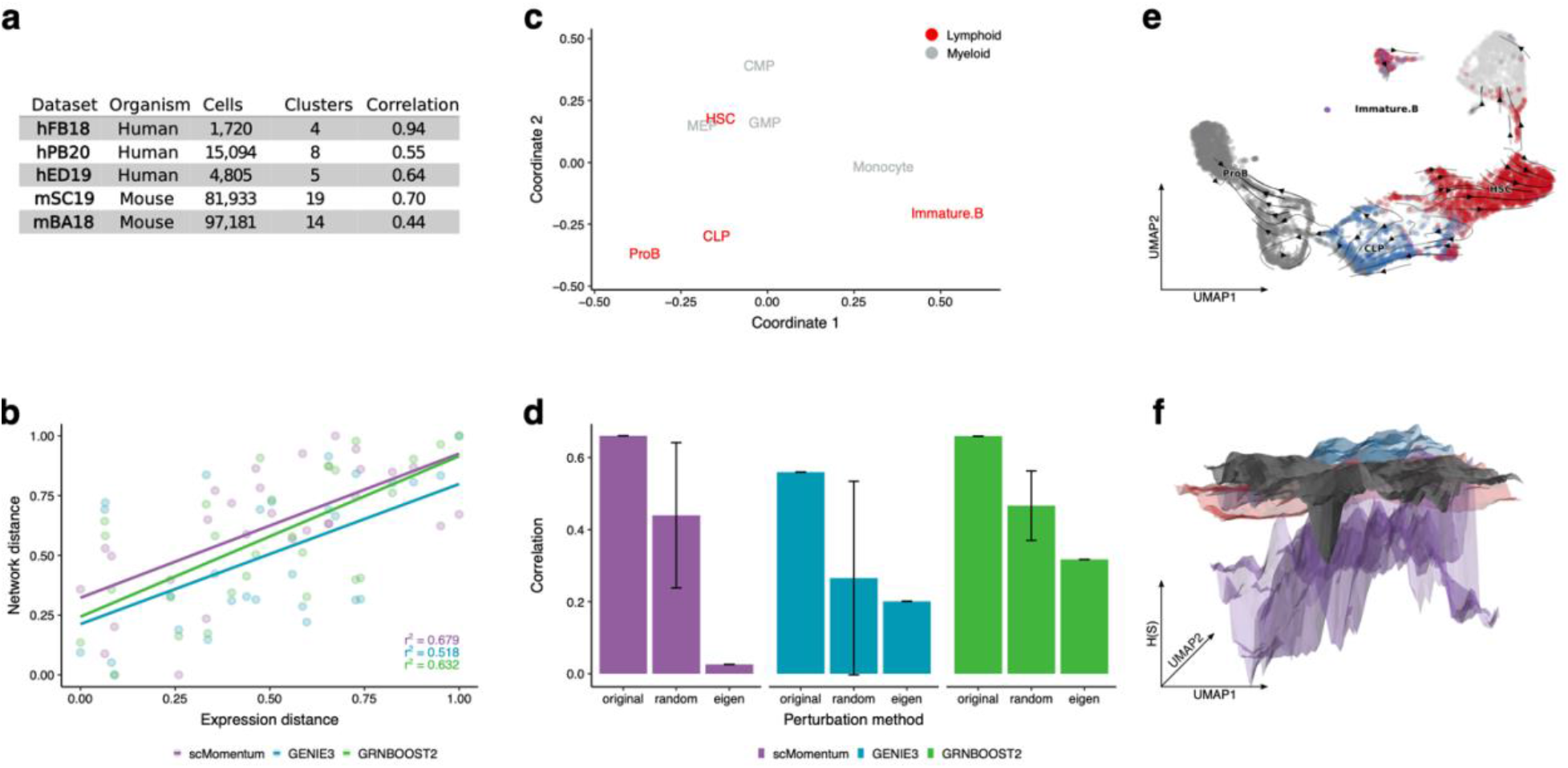
scMomentum recovers relevant biological features of cell types in multiple real datasets. **(a)** Mantel correlations of distance matrices were derived for five real datasets (description in Methods); all correlations were significant at a p-value cut-off of 0.05. **(b)** Methods’ benchmark based on the preservation of cluster distances; see Fig 1e caption for a detailed description; all correlations were significant at a p-value cut-off of 0.05. **(c)** MDS map inferred from the network-derived distance matrix. **(d)** Mantel correlations for each inference method under each perturbation setting (shown in the x-axis); for the method “random” error bars show the standard deviation of 10 independent perturbations). **(e)** Cell-velocity map of the mBD20 dataset; annotations are shown for cell types belonging to the Lymphoid lineage (HSC, CLP, ProB, and Immature B), the remaining cells in the dataset are colored in gray. **(f)** Energy landscape of cell types from the Lymphoid lineage; Surface colors correspond to the annotations in (e).

To assess our networks’ ability to capture cellular dynamics, we looked at cellular differentiation and response to targeted perturbations. The network distance matrix recovered trajectories on a Multidimensional Scaling projection (MDS) that resemble cell progressions along hematopoiesis (Figure 2c), suggesting that the networks capture cell-type-specific properties underlying their developmental progressions. Moreover, this result was robust to cellular noise and information loss (see Supplementary material). Then, we took a gene-centered approach and removed 30% of the genes with the largest eigenvalue’s centralities (see Methods) and found that the correlation between distance matrices was lower than that of a random perturbation (Figure 2d). This result shows that our networks detected a set of essential network regulators, a common feature of biological networks. This was not the case for other methods.

To investigate the developmental potential captured by our networks, we reconstructed an analog of the Waddington landscape for the lymphoid lineage (Figure 2e and f). Interestingly, HSC and CLP landscapes have the highest energy and lack steep regions, highlighting their pluripotency and tendency to continue differentiating. The Pro-B cell landscape has a deep basin that contains the vast majority of cells (see Supplementary material), suggesting that they are “trapped” at this stage and might only progress along a confined path. The next stage is Immature B cells, which have the lowest energy and are positioned right underneath the basin of Pro-B cells, showing a possible direction of differentiation. Thus, our networks allow the reconstruction of energy landscapes capturing the corresponding cells’ biology [17].

A significant advantage of our approach is the extraction of weights and directions within the networks. To further assess its biological relevance, we computed an activator/repressor score by adding up all the positive/negative outgoing edges of every gene in the cells along both hematopoietic lineages (Figure 3). Notably, we analyzed Prothymosin Alpha (PTMA), a proteincoding gene involved in immune function modulation [18]. The expression and velocity of PTMA follow similar trends in both lineages. Although, it has a strong activator score in CLPs and a strong repressor score in GMPs. This observation raises the intriguing possibility that some of the expression changes previously associated with alterations of PTMA [19] might be the effect of altered cellular development, rather than expression changes alone. The inferred weights are also useful to discover genes with dynamic properties within the network, providing insights on possible cell reprogramming undetectable by gene expression or velocity alone.

**Figure 3.**
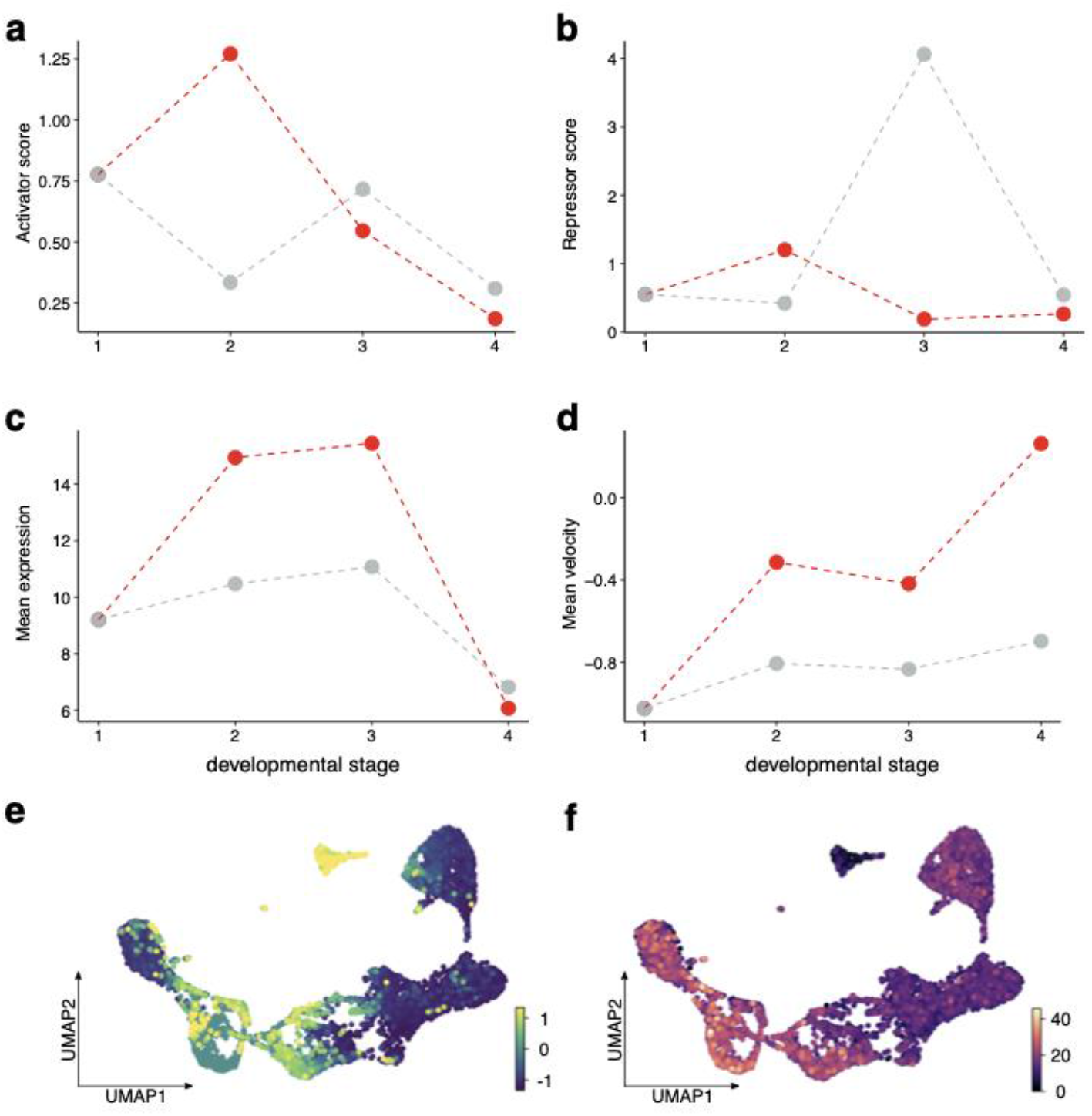
scMomentum uncovers the dynamics of transcriptional regulation of PTMA during hematopoiesis. Panels **a-d** order in the x-axis corresponds to the cell-types order along each lineage. Both start with HSC; the progression in the Lymphoid lineage is (2) CLP, (3) ProB, (4) Immature B; and in the myeloid lineage is (2) CMP, (3) GMP, (4) Monocyte. **e-f** shows the UMAP embedding containing all the cells in the dataset. In **(e),** each cell is colored by the corresponding value of velocity for PTMA only. In **(f),** the color scale represents PTMA’s expression.

Here we showed that incorporating RNA velocity into the inference of cell-type-specific GRNs allows us to model the regulatory mechanisms underlying dynamic developmental processes. The resulting networks harbor cell-type-specific regulatory properties that made it possible to reconstruct an interpretable Waddington landscape analog. Conceptually, these networks capture different states corresponding to different cell types, each with *momentum* to move in a landscape. This interpretation, constructed from the information within GRNs, opens up the possibility to study how regulatory processes shape cellular development in multiple contexts [20]. Moreover, scMomentum has a significant computational advantage over previous approaches, which would facilitate its application to the vast compendium of existing scRNA-seq datasets and open the prospect of building a Cell Regulatory Atlas. To our knowledge, this is the first cell-type-specific GRN inference method that scales to large datasets and recovers directed, signed, and weighted GRNs in a data-driven manner.

## Acknowledgments

We acknowledge B. Li and J. Ye for initial guidance and discussion of the mathematical model. We also thank J. Ye for helping with the annotations of one of the public datasets. F.P acknowledges funding obtained Instituto de Salud Carlos II (PI17/00701, PI20/01308 and CB16/12/00489) co-funded by FEDER grant, and the AGATA grant (0011-1411-2020-000011 and 0011-1411-2020-000010) from the Government of Navarra. NAK was supported by the Karolinska Institute’s funds and KA Wallenberg Foundation (KAW 2017.0077). L.M.S and J.P.B-T were supported by a VSRP fellowship from King Abdullah University of Science and Technology.

## Competing interests

Authors declare no competing interests.

## Materials and Methods

### Network inference

We derive networks in a cell-type-specific manner. Therefore, the method can readily be combined with a user-defined choice of appropriate clustering and annotation pipelines to process the data. For specific pre-processing details, see Supplementary Methods. For each cell type, we re-define the change in gene expression over time as the contribution from all the other genes expressed in the cluster have on its mRNA expression and the level of degradation of its mRNA as follows:

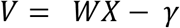

Where *X* is the cell-by-gene expression matrix, and *V* is the cell-by-gene velocity matrix. The diagonal matrix *γ* contains gene-specific degradation constants (both *V* and *γ* are retrieved from scVelo). *W* is the gene by gene weighted and directed adjacency matrix (inferred GRN). Then we solve for *W*, obtaining the final model:

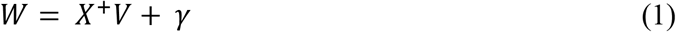

Since we solve Eqn. 1 as an overdetermined system using least squares, the network’s size is bounded by the number of cells in each cluster.

### Calculation of distances between clusters and between networks

For every cluster in the data set, we calculate the Euclidean distance with all the remaining clusters. In the expression-derived mode, for every pair of clusters, the distance between them is defined as:

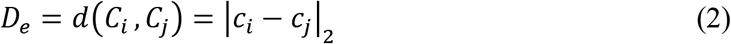

Where *c_i_* and *c_j_* are the mean expression vectors of clusters *C_i_* and *C_j_*, respectively. This is used as a reference distance matrix.

In the network-derived mode, we define the distance between every pair of clusters as:

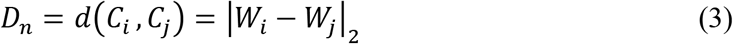

Where *W_i_* and *W_j_* denote the adjacency matrices derived from clusters *C_i_* and *C_j_*, respectively. To test the accuracy of the inferred networks, we calculate

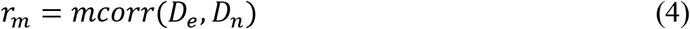

Where *mcorr* refers to the Mantel correlation of distance matrices, which accounts for the inherent symmetry and row-column relationship of *D*. Although *W_i_* and *W_j_* have the same dimensions; they might not have the same genes. To accommodate this when computing the correlations, we use the set of genes *G* in each network to find *G_i_* ∪ *G_j_*, the universal set of genes *G_u_*. The uniform distribution is used to sample the entries of *W_i_* corresponding to 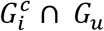 and those of *W_j_* corresponding to 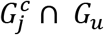.

### Selection of genes

A critical step in any single-cell analysis is choosing the appropriate set of genes. We tested six different approaches to rank and select varying numbers of genes within each cluster, and *r_m_* (Eqn. 4) to rank them. The ranking schemes were based on absolute gene velocity, signed gene velocity, velocity variance, expression and expression variance. We tested their combination with different network sizes, ranging from 50 to 500 genes in steps of 50. In each data set, we selected the combination of ranking and sized with the highest, *r_m_* value.

### Landscape reconstruction

For every network *W_i_*, an energy landscape is reconstructed using the Discrete Hopfield Network (DHN) formalism [1]. A DHN is formed by *n* neurons 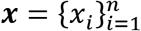 that can be in two different states ON or OFF. At each discrete time step, a subset of these neurons change state depending on the influence of all other neurons weighted by the interaction network *W_i_*:

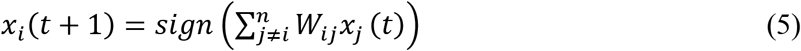

In the original paper [1] Hopfield demonstrated the existence of an energy function such that

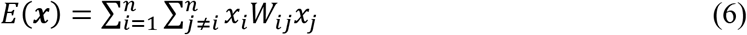

If the interaction matrix *W_ij_* is symmetric, after evolving the system according to Eqn. 5 the system would move to the states corresponding to local minima of the function.

The original bottom-up idea underlying the DHN is that the *W* matrix can be constructed so that it stores certain fates, corresponding to local minima of the energy function to which the system would evolve.

Our top-down approach corresponds to analyzing the structure of the energy landscape after taking the GRN as interaction matrix for a DHN where each gene corresponds to a neuron. To analyze the energy of a cell, its expression must first be discretized to the ON and OFF states. To accomplish this is to consider a gene to be ON or OFF depending if it is above or below a certain predefined threshold for each gene. This threshold can be taken as the mean value of that gene over all the cells, the median of the gene expression over all the cells, or other values that might be biologically important.

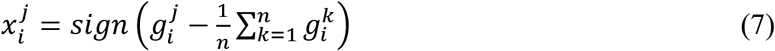

The landscape is displayed over a two-dimensional space for visualization purposes. Therefore, we use PCA or any embedding algorithm where an inverse exists for the data over two dimensions.

First, the two-dimensional embedding, 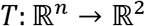, is calculated and used to project the cells. A square containing all the projected cells is gridded into *N* × *M* grid-points, 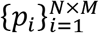, which are then pushed back to the high-dimensional space with the inverse of the embedding, 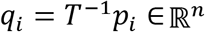. These points are then discretized as in equation (7), and the energy for each point is calculated. In this way, the energy of each grid-point is calculated and can be plot as a surface over the 2d space containing the projections of the cells, for which the energy is also calculated and plot on top of the grid.

### Network perturbations

For every network *W_i_*, we calculate a vector of eigenvalues *λ_i_*, and remove from *W_i_* all the entries of the genes with the top 30% eigenvalues in *λ_i_*. Then, we estimate the distance *D_n,p_* between the perturbed networks to calculate *mcorr*(*D_e_, D_n,p_*), and use *r_m,p_* as a measure of network robustness.

### Gene-specific network scores

For a gene *g_k_* in a network *W_i_* where *k* goes from 1,…, *N_i_* and *N_i_* is the number of genes in *W_i_*, we compute an activator score *act* as

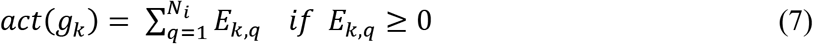

Where *E_k,q_* refers to the outgoing edge from *g_k_* to *g_q_*. Similarly, we calculate a repressor score *rep* as

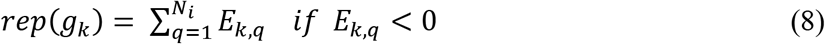

### Benchmark framework

We defined the best-performing set of genes as previously described to infer cell-type-specific manner using GENIE3 [2], GRNBOOST2 [3] and PIDC [4] in the simulated and the in-house generate data sets. We discarded PIDC since often the clusters did not contain enough cells to meet its data requirements. We benchmarked the methods using the set of *r_m_* values estimated by comparing the networks derived from each method to the same *D_e_* matrix.

### Data

#### Simulated

We simulated a dataset with Dyngen v0.4.0 [5], using *backbone_disconnected* with left and right backbones set to *backbone_consecutive_bifurcating*. Models were initialized with the following parameters: *num cells 20.000, num_tfs 50, num targets 200, num_hks 250*.

#### Public

##### Human fetal forebrain (hFB18)

Human fetal forebrain cells from 10-week fetal tissue were generated in *La Manno et al*., 2018. This dataset is accessible from the SRA under the accession code SRP129388 [6].

##### Human PBMCs (hPB20)

10X Genomics 5K PBMC dataset downloaded from the company’s website [7].

##### Mouse developing spinal cord (mSC19)

Mouse embryos from stages E9.5 to E13.5 from *Delile et al., 2019*. Raw sequencing files were retrieved from the database ArrayExpress under the accession E-MTAB-7320 [8].

##### Mouse brain atlas (mBA18)

Adult mouse nervous system data set generated by *Zeisel et al., 2018*. It is deposited in the SRA under the accession code SRP135960 [9].

##### Human embryonic hematopoiesis (hED19)

Human embryonic sections were collected from the Carnegie at stage 12 to 14. The data generated in *Zeng et al., 2019* is available at NCBI’s Gene Expression Omnibus (GEO) with the accession code GSE135202 [10].

##### Human hematopoiesis - In house (mBD20)

Cell sorting was performed using a FACSAria (BD Biosciences) and analyzed with FACSdiva software (BD Biosciences). Standard, strict forward scatter width versus area criteria were used to discriminate doublets and gate only singleton cells. Viable cells were identified by staining with 7-AAD (BD Bioscience). HSC cells were extracted from the bone marrow using CD34+ membrane marker. According to the manufacturer’s instructions, the transcriptome of the cells was profiled using Single Cell 3’ Reagent Kits v3 (10X Genomics).

Sequenced libraries were demultiplexed, aligned to the human transcriptome (GRC3h8/hg20) and quantified using Cell Ranger (3.0.1) from 10X Genomics. The output of the pre-processing pipeline consisted of UMI-derived expression matrices per cell. Quality control filters applied for filtering the cells were: the number of detected genes, the number of UMIs, and the proportion of UMIs mapped to mitochondrial genes per cell. The thresholds for each of the single-cell libraries were selected based on the distribution of the variables enumerated. Count-based matrices were subjected to normalization, identification of highly variable genes, and removal of unwanted sources of variation using Seurat3 [11] Next, cells were labeled to the different cell populations shown using SingleR [12]. The annotation was conducted using as a reference an In-house bulk RNA-seq from the enumerated populations.

## Notes

https://github.com/larisa-msoto/scMomentum.git

### Competing Interest Statement

The authors have declared no competing interest.

## References

1. Han, X. et al. Construction of a human cell landscape at single-cell level. Nature 581, 303–309 (2020).

2. Rozenblatt-Rosen, O., Stubbington, M. J. T., Regev, A. & Teichmann, S. A. The Human Cell Atlas: from vision to reality. Nature 550, 451–453 (2017).

3. Marbach, D. et al. Wisdom of crowds for robust gene network inference. Nature Methods 9, 796–804 (2012).

4. Pratapa, A., Jalihal, A. P., Law, J. N., Bharadwaj, A. & Murali, T. M. Benchmarking algorithms for gene regulatory network inference from single-cell transcriptomic data. Nature Methods 17, 147–154 (2020).

5. Kharchenko, P. V., Silberstein, L. & Scadden, D. T. Bayesian approach to single-cell differential expression analysis. Nature Methods 11, 740–742 (2014).

6. Waddington, C. H. The Strategy of the Genes: A Discussion of some Aspects of theoretical Biology. (George Allen & Unwin, 1957).

7. Matsumoto, H. et al. SCODE: an efficient regulatory network inference algorithm from single-cell RNA-Seq during differentiation. Bioinformatics 33, 2314–2321 (2017).

8. Gao, N. P., Ud-Dean, S. M. M., Gandrillon, O. & Gunawan, R. SINCERITIES: inferring gene regulatory networks from time-stamped single cell transcriptional expression profiles. Bioinformatics 34, 258–266 (2017).

9. Specht, A. T. & Li, J. LEAP: constructing gene co-expression networks for single-cell RNA-sequencing data using pseudotime ordering. Bioinformatics (2016). doi:10.1093/bioinformatics/btw729

10. Morgan, D. et al. Perturbation-based gene regulatory network inference to unravel oncogenic mechanisms. Scientific Reports 10, (2020).

11. Tegnér, J. & Björkegren, J. Perturbations to uncover gene networks. Trends in Genetics 23, 34–41 (2007).

12. Manno, G. L. et al. RNA velocity in single cells. (2017). doi:10.1101/206052

13. Hopfield, J. J. Neurons with graded response have collective computational properties like those of two-state neurons. Proceedings of the National Academy of Sciences 81, 3088–3092 (1984).

14. Cannoodt, R., Saelens, W., Deconinck, L. & Saeys, Y. dyngen: a multi-modal simulator for spearheading new single-cell omics analyses. (2020). doi:10.1101/2020.02.06.936971

15. Huynh-Thu, V. A., Irrthum, A., Wehenkel, L. & Geurts, P. Inferring Regulatory Networks from Expression Data Using Tree-Based Methods. PLoS ONE 5, (2010).

16. Moerman, T. et al. GRNBoost2 and Arboreto: efficient and scalable inference of gene regulatory networks. Bioinformatics 35, 2159–2161 (2018).

17. Kondo, M. Lymphoid and myeloid lineage commitment in multipotent hematopoietic progenitors. Immunological Reviews 238, 37–46 (2010).

18. Samara, P., Ioannou, K. & Tsitsilonis, O. Prothymosin Alpha and Immune Responses. Vitamins and Hormones Thymosins 179–207 (2016). doi:10.1016/bs.vh.2016.04.008

19. Kobayashi, T. et al. Overexpression of the Oncoprotein Prothymosin α Triggers a p53 Response that Involves p53 Acetylation. Cancer Research 66, 3137–3144 (2006).

20. Tegnér, J. & Björkegren, J. Perturbations to uncover gene networks. Trends in Genetics 23, 34–41 (2007).

## References

1. Hopfield, J. J. Neural Networks and Physical Systems with Emergent Collective Computational Abilities. Feynman and Computation 7–19 doi:10.1201/9780429500459-2

2. Huynh-Thu, V. A., Irrthum, A., Wehenkel, L. & Geurts, P. Inferring Regulatory Networks from Expression Data Using Tree-Based Methods. PLoS ONE 5, (2010).

3. Moerman, T. et al. GRNBoost2 and Arboreto: efficient and scalable inference of gene regulatory networks. Bioinformatics 35, 2159–2161 (2018).

4. Chan, T. E., Stumpf, M. P. & Babtie, A. C. Gene Regulatory Network Inference from Single-Cell Data Using Multivariate Information Measures. Cell Systems 5, (2017).

5. Cannoodt, R., Saelens, W., Deconinck, L. & Saeys, Y. dyngen: a multi-modal simulator for spearheading new single-cell omics analyses. (2020). doi:10.1101/2020.02.06.936971

6. Manno, G. L. et al. RNA velocity of single cells. Nature 560, 494–498 (2018).

7. Datasets. 10x Genomics (2019). Available at: http://www.10xgenomics.com/resources/datasets/.

8. Delile, J. et al. Single cell transcriptomics reveals spatial and temporal dynamics of gene expression in the developing mouse spinal cord. Development 146, (2019).

9. Zeisel, A. et al. Molecular Architecture of the Mouse Nervous System. Cell 174, (2018).

10. Zeng, Y. et al. Tracing the first hematopoietic stem cell generation in human embryo by single-cell RNA sequencing. Cell Research 29, 881–894 (2019).

11. Stuart, T. et al. Comprehensive Integration of Single-Cell Data. Cell 177, (2019).

12. Aran, D. et al. Reference-based analysis of lung single-cell sequencing reveals a transitional profibrotic macrophage. Nature Immunology 20, 163–172 (2019).

